# The Transcription Factor TEAD4 Enhances Lung Adenocarcinoma Malignancy through Enhancing PKM2 Mediated Glycolysis

**DOI:** 10.1101/2020.11.25.399295

**Authors:** Yan Hu, Hanshuo Mu, Zhiping Deng

## Abstract

Lung adenocarcinoma (LUAD) is a deadly disease with a hallmark of aberrant metabolism. The role of TEAD4 in LUAD is rarely reported as well as its potential mechanisms. The expression level of TEAD4 and PKM2 was measured by quantitative real-time PCR and western blot. Luciferase reporter assay were employed to verify the effect of TEAD4 on PKM2 promoter as well as TEAD4/PKM2 axis on reporter activity of HIF-1α. Glycolysis was investigated according to glucose consumption, lactate production and the extracellular acidification rate (ECAR). Cell Counting Kit-8 was used to assess cell viability. The present study indicated that TEAD4 and PKM2 were upregulated in LUAD and closely related to prognosis. Mechanistic investigations identified that TEAD4 played a key role as a transcription factor and promoted PKM2 transcription and expression, which further altered the reporter activity of HIF-1α and upregulated HIF-1α-targeted glycolytic genes GLUT1 and HK2. Functional assays revealed that TEAD4 and PKM2 affected glycolytic and 2-DG blocked the positive function of TEAD4 and PKM2 on glycolytic. Besides, TEAD4/PKM2 axis affects LUAD cells survival through glycolysis. Together, these data provided evidence that both TEAD4 and PKM2 were poor prognosticator. Targeting TEAD4/PKM2 axis might be an effective therapeutic strategy for LUAD.

## Introduction

Lung cancer is the most diagnosed cancer and the leading cause of cancer-related deaths in United States with 142,670 death in 2019 Siegel et al., 2019. Lung cancer is mainly composed of small cell lung cancer (SCLC) and non-small cell lung cancer (NSCLC) Brambilla et al., 2001. Lung adenocarcinoma (LUAD) is the most general subtype of NSCLC, accounting for 50% of the cases Zappa and Mousa, 2016. At present, LUAD is primarily treated by surgery, assisted with radiotherapy and chemotherapy, but LUAD is still one of the most aggressive and deadly cancers, with poor outcomes of patients Denisenko et al., 2018. Hence, it is important to understand the underlying mechanisms that regulate LUAD pathogenesis and explore new targets for the treatment of LUAD.

TEA domain (TEAD) proteins are a family of transcription factors that adjust the transcription of various genes related to cell proliferation, apoptosis and metastasis Shi et al., 2017. There are four members in TEAD family, termed TEAD1~4. TEAD proteins were reported to bind the simian virus 40 (SV40) enhancer and activate downstream transcription Xiao et al., 1987, Davidson et al., 1988. Recently, the pivotal role of TEAD proteins in the development of various malignancies, such as breast, ovarian prostate and liver cancers has been clearly confirmed Zhou et al., 2016. TEAD4, a member of TEAD family, has been identified as a potential therapeutic target and prognostic marker in both breast cancer and gastric cancer Chan et al., 2009, Lim et al., 2014. However, the role of TEAD4 in NSCLC is rarely reported and the purpose in this study is to explore the effect of TEAD4 in LUAD as well as its potential mechanisms.

Our results demonstrate that TEAD4 and Pyruvate kinase isozymes M2 (PKM2) were highly expressed in LUAD tissues and could serve as prognostic marker in LUAD. PKM2, which played an important role in glycolytic and cancer progression Hua et al., 2020, Chen et al., 2020, was recognized to be a downstream effector of TEAD4 in LUAD. Our work identified TEAD4 as a regulator of glycolytic, which may be a promising metabolism blocker target for antitumor therapy.

## Results

### TEAD4 and PKM2 are up-regulated patients with LUAD

We firstly detected the transcription level of TEAD4 by qRT☐PCR in a total of 20 pairs of clinical LUAD tissues and adjacent healthy tissues. The mRNA levels of TEAD4 in LUAD tissues were significantly increased when compared with the adjacent one (Fig. 1A). To further explore the clinical significance of TEAD4 in NSCLC, we evaluated its prognostic effect via a public database of Kaplan☐Meier plotter analysis (http://www.kmplot.com). Higher TEAD4 level was correlated with poorer overall survival (OS) in LUAD while there seemed no predictive significance of TEAD4 in patients with Lung squamous cell carcinoma (LUSC) (Fig. 1B-C). These results revealed a hopeful role of TEAD4 in help predicting clinical outcome of patients with LUAD. Similarly, PKM2 also was found to be upregulated in LUAD tissues and high expression of PKM2 was significantly associated with a poor prognosis in these patients with LUAD (Fig. 1D-F). Considering the tumor☐promoting role of TEAD4 and PKM2 in LUAD, we investigated the mRNA and protein levels of TEAD and PKM2 in 2 human LUAD cell lines (A549 and NCI☐H1299) and one normal bronchial epithelial cell line (HBE) as control. The results suggested that TEAD and PKM2 levels were notably amplified in LUAD cell lines (Fig. 1G-H). In short, both TEAD and PKM2 could be viewed as potential biomarker in LUAD.

**Figure 1.**
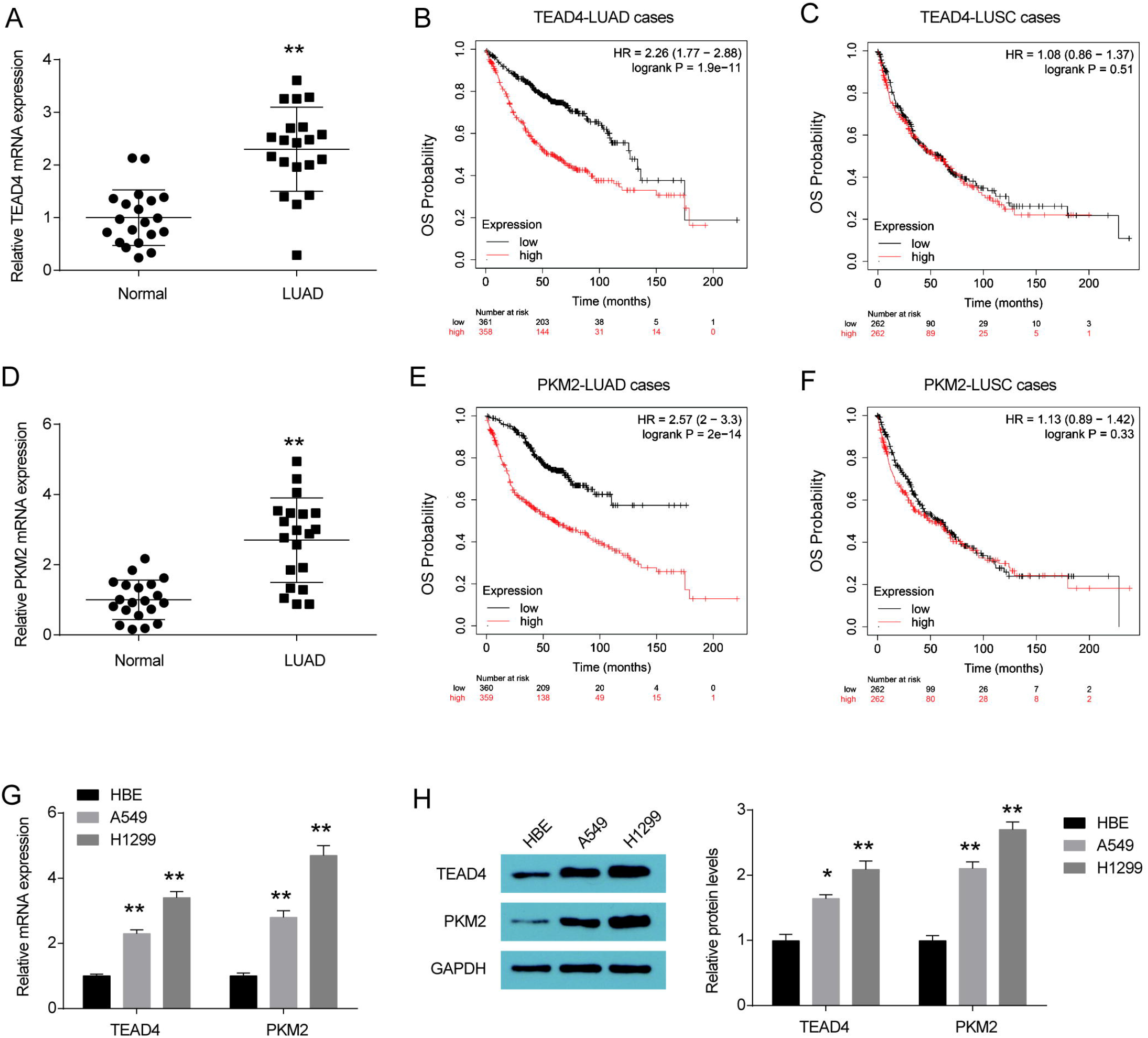
TEAD4 and PKM2 are clinically relevant in LUAD. (A) TEAD4 mRNA levels in lung adenocarcinoma (LUAD) tissues and matched adjacent healthy tissues (n = 20). (B) The overall survival (OS) was analyzed and compared between patients with low and high levels of TEAD4 in LUAD patients from database of Kaplan-Meier plotter. (C) The OS was analyzed and compared between patients with low and high levels of TEAD4 in LUSC patients from database of Kaplan-Meier plotter. (D) TEAD4 mRNA levels in LUAD tissues and matched adjacent healthy tissues (n = 20). (E) The OS of PKM2 in LUAD patients were analyzed by Kaplan-Meier plotter. (F) The OS of PKM2 in LUSC patients were analyzed by Kaplan-Meier plotter. (G) TEAD4 and PKM2 mRNA levels in human LUAD cell lines and normal bronchial epithelial cell line (HBE). (H) TEAD4 and PKM2 protein levels in human LUAD cell lines and normal bronchial epithelial cell line (HBE). \**P*<0.05, \*\**P*<0.01 vs. Normal tissues or HBE cells.

### TEAD4 promotes transcriptional activation of PKM2

We next explored the regulatory mechanism between TEAD4 and PKM2, the results form database of GEPIA (http://gepia.cancer-pku.cn/index.html) indicated a positive correlation between TEAD4 and PKM2 (Fig. 2A). TEAD4 overexpression signally enhanced TEAD4 and PKM2 mRNA levels while TEAD4 knockdown through specific short interfering RNA (siRNA) clearly inhibited TEAD4 and PKM2 mRNA levels in A549 and NCI☐H1299 cells (Fig. 2B-C). On the contrary, both PKM2 overexpression and knockdown showed no significance change on TEAD4 mRNA levels in A549 and NCI☐H1299 cells (Fig. 2D-E). These results revealed that TEAD4 was an upstream regulator of PKM2. Transcription factor TEAD4 was reported to be an upstream regulator in many human cancers Zhang et al., 2018, Shuai et al., 2020, He et al., 2019. We next analyzed ChIP-Seq data of A549 downloaded from the Encyclopedia of DNA Elements (ENCODE) database and found that TEAD4 was highly enriched in the PKM2 promoter regions (Fig. 2F). To further understand the effects of TEAD4 on PKM2 transcription, we subcloned promoter sequence of PKM2 into p-GL3-Basic vectors and transfected them to A549 and H1299 cells. The firefly luciferase activity in p-GL3-promoter group was higher than p-GL3-Basic group. More importantly, TEAD4 overexpression markedly enhanced PKM2 promoter activity and TEAD4 knockdown reduced PKM2 promoter activity in A549 and H1299 cells (Fig. 2G-H). Finally, TEAD4 overexpression signally upregulated TEAD4 and PKM2 protein levels while TEAD4 knockdown inhibited TEAD4 and PKM2 protein levels (Fig. 2I). In a word, TEAD4 serve as a transcription factor which is highly enriched in the PKM2 promoter regions and promotes transcriptional activation of PKM2.

**Figure 2.**
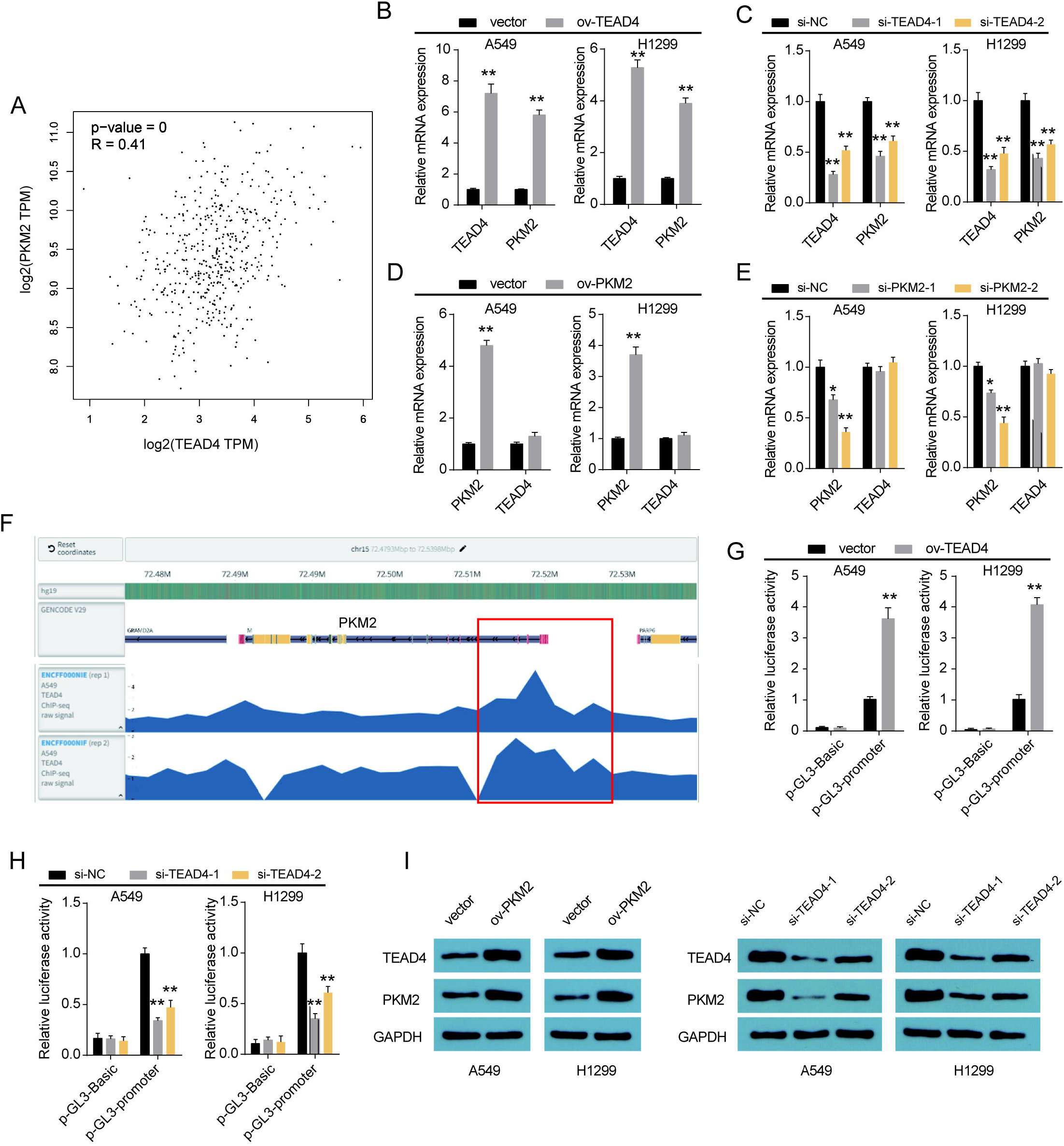
TEAD4 promotes PKM2 transcription. (A) TEAD4 and PKM2 levels in LUAD tissues from database of GEPIA showed a positive correlation. (B) TEAD4 and PKM2 mRNA levels in A549 and NCI☐H1299 cells after TEAD4 overexpression. \*\**P*<0.01 vs. vector. (C) TEAD4 and PKM2 mRNA levels in A549 and NCI☐H1299 cells after TEAD4 knockdown. \*\**P*<0.01 vs. si-NC. (D) TEAD4 and PKM2 mRNA levels in A549 and NCI☐H1299 cells after PKM2 overexpression. \*\**P*<0.01 vs. vector. (E) TEAD4 and PKM2 mRNA levels in A549 and NCI☐H1299 cells after PKM2 knockdown. \*\**P*<0.01 vs. si-NC. (F) Analysis of TEAD4 ChIP-seq of A549 cells in the PKM2 locus. (G) Luciferase activity in A549 and NCI☐H1299 cells transfected with luciferase reporter pGL3-promotor or TEAD4 overexpression plasmid. Data are shown as the relative ratio of firefly luciferase activity to Renilla luciferase activity. \*\**P* <0.01 vs. vector. (H) Luciferase activity in A549 and NCI☐H1299 cells transfected with luciferase reporter pGL3-promotor or TEAD4 siRNAs. \*\**P* <0.01 vs. si-NC. (I) TEAD4 and PKM2 protein levels in A549 and NCI☐H1299 cells after TEAD4 overexpression or knockdown.

### TEAD4 and PKM2 are induced under metabolic stress and play a crucial role on glycolytic metabolism reprogramming

As is well-known that glucose deprivation is a widespread characteristic of solid tumors and PKM2 catalyzes irreversible committed step of glycolysis Hua et al., 2020, Hou et al., 2020. We further explored whether TEAD4 and PKM2 played a role in glycolytic metabolism under metabolic stress. Different glucose concentrations from 2.5 mM to 25 mM were employed to simulate glucose deprivation conditions. TEAD4 and PKM2 mRNA levels were escalated along with the decrease glucose concentrations (Fig 3A). The 2-DG was utilized to block glucose supply. As shown in Figure 3B, 2-DG promoted TEAD4 and PKM2 mRNA levels in a dose-dependent, indicating that TEAD4/PKM2 axis might hold a vital effect in cell adaptation to metabolic stress. TEAD4 overexpression increased glucose uptake, lactate secretion, and ECAR whereas PKM2 knockdown hold a opposite effect in A549 and H1299 cells as compared to control group. PKM2 knockdown reversed the promotion of glycolytic induced by TEAD4 overexpression (Fig 3C-F). In short, TEAD4/PKM2 axis were involved in metabolic stress and participated in glycolytic metabolism reprogramming.

**Figure 3.**
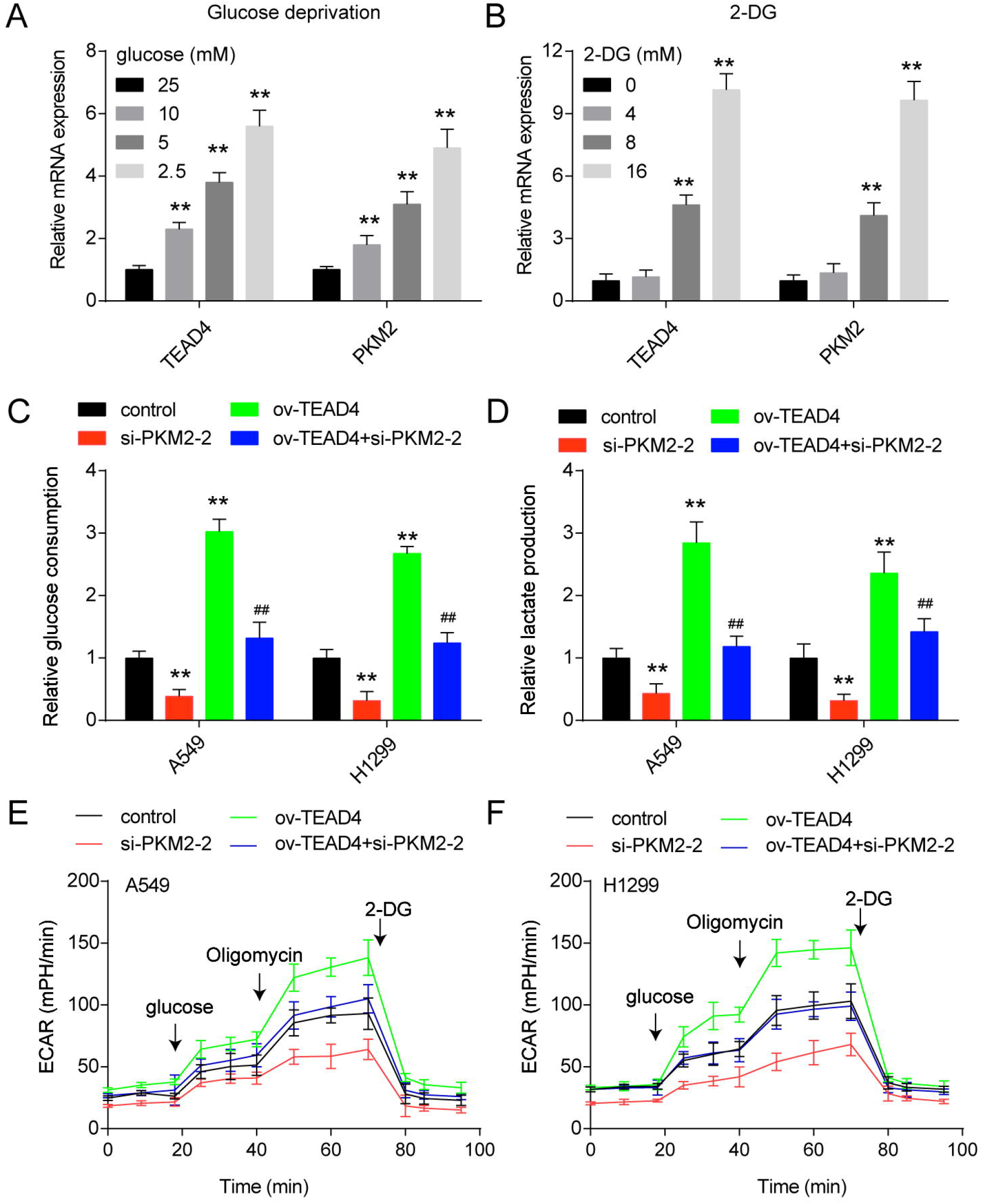
TEAD4/PKM2 axis plays a crucial role on glycolytic metabolism reprogramming. (A) TEAD4 and PKM2 mRNA levels in A549 after treated with different glucose concentrations from 2.5 mM to 25 mM. \*\**P*<0.01 vs. 25 mM. (B) TEAD4 and PKM2 mRNA levels in A549 after treated with different 2-DG concentrations from 0 mM to 16 mM. \*\**P*<0.01 vs. 0 mM. (C) Detection of glucose consumption in A549 and NCI☐H1299 cells were detected after transfection for 48 h. \*\**P*<0.01 vs. control, ##*P*<0.01 vs. ov-TEAD4. (D) Detection of lactate excretion in A549 and NCI☐H1299 cells were detected after transfection for 48 h. \*\**P*<0.01 vs. control, ##*P*<0.01 vs. ov-TEAD4. (D) Detection of glycolytic capacity (ECAR) in A549 and NCI☐H1299 cells were detected after transfection for 48 h.

### TEAD4/PKM2 axis actives HIF-1α and promotes glucose transporter and metabolic enzymes expression

PKM2 is known to play a pivotal role in glycolysis through activating HIF-1α-dependent transcription of glycolytic enzymes Luo et al., 2011. TEAD4 overexpression substantially enhanced the reporter activity of HIF-1α and PKM2 knockdown markedly suppressed the reporter activity of HIF-1α in A549 and H1299 cells. PKM2 knockdown reversed the promotion induced by TEAD4 overexpression (Fig 4A). TEAD4 overexpression upregulated HIF-1α protein levels and PKM2 knockdown inhibited HIF-1α protein levels (Fig 4B-E). TEAD4/PKM2 axis also was found to change the expression of HIF-1α-targeted glycolytic genes, such as glucose transporter-1 (GLUT1) and hexokinase II (HK2) (Fig 4B-E). It is likely that TEAD4/PKM2 axis remodel glucose metabolism in LUAD cells via activating HIF-1α.

**Figure 4.**
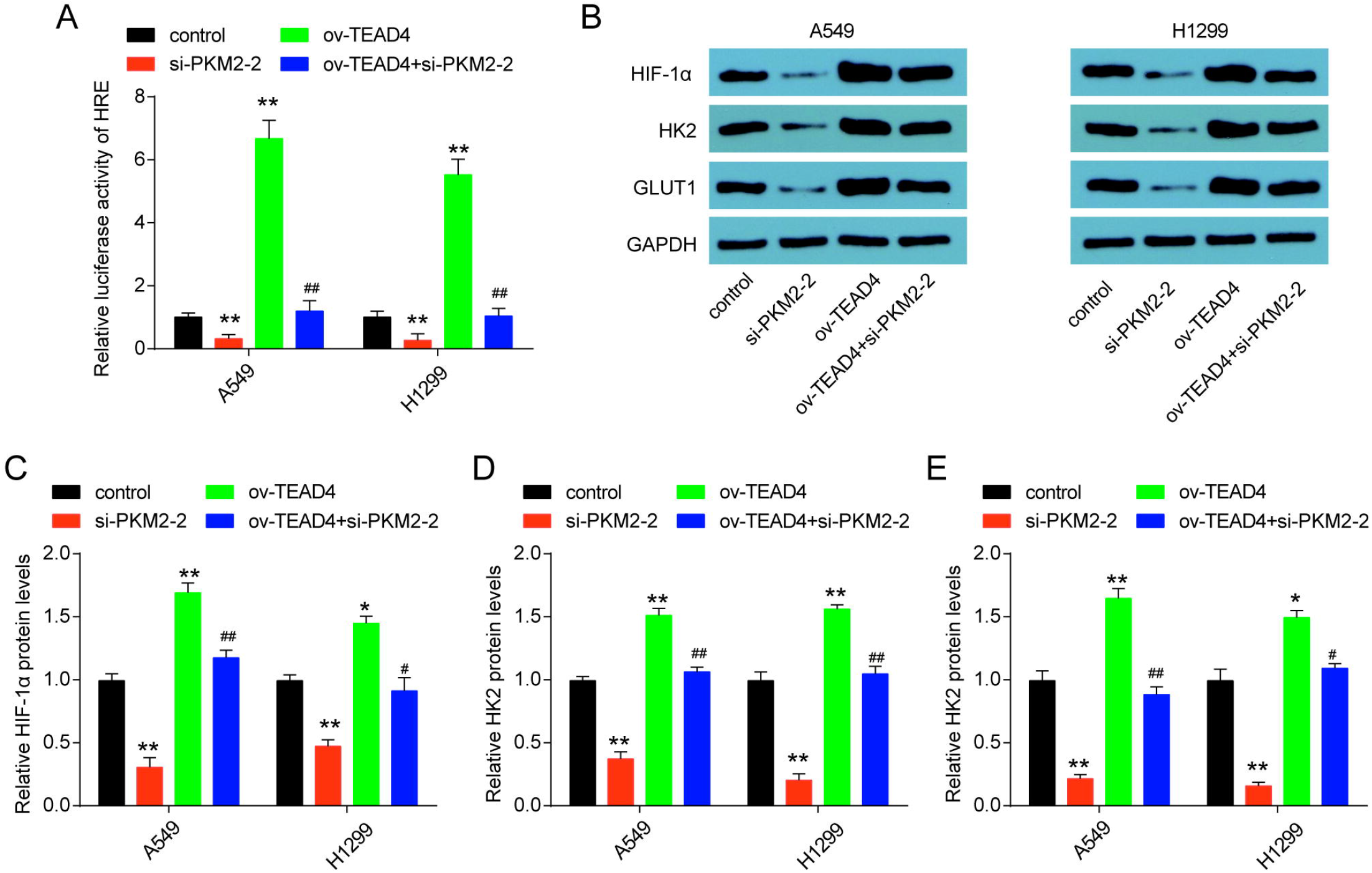
TEAD4/PKM2 axis actives HIF-1α and promotes GLUT1 and HK2 expression. (A) HIF-1α responsive luciferase reporter were examined after indicated transfection for 48 h in A549 and NCI☐H1299 cells. (B) HIF-1α, HK2 and GLUT1 protein levels were examined after indicated transfection for 48 h in A549 and NCI☐H1299 cells. (C-E) The quantification of HIF-1α, HK2 and GLUT1 protein levels in A549 and NCI☐H1299 cells. \**P*<0.05, \*\**P*<0.01 vs. control; #*P*<0.05, ##*P*<0.01 vs. ov-TEAD4.

### TEAD4/PKM2 axis affects LUAD cells survival through glycolysis

Uncontrolled, unlimited and accelerated multiplication is one of the most fundamental biologic behavior of cancer cells. To further explored whether TEAD4/PKM2 axis affects LUAD cells survival through glycolysis, glycolytic inhibitor 2-DG was employed. We found that TEAD4 overexpression or PKM2 overexpression promoted A549 and H1299 cell survival. However, glycolytic inhibitor 2-DG observably inhibited A549 and H1299 cell survival. These effects were completely reversed by the 2-DG (Fig 5A-B). What’s more, TEAD4/PKM2 axis also was found to control A549 and NCI☐H1299 cell survival (Fig 5C). These results suggested that TEAD4/PKM2 axis was involved in malignant proliferation of LUAD via glycolysis.

**Figure 5.**
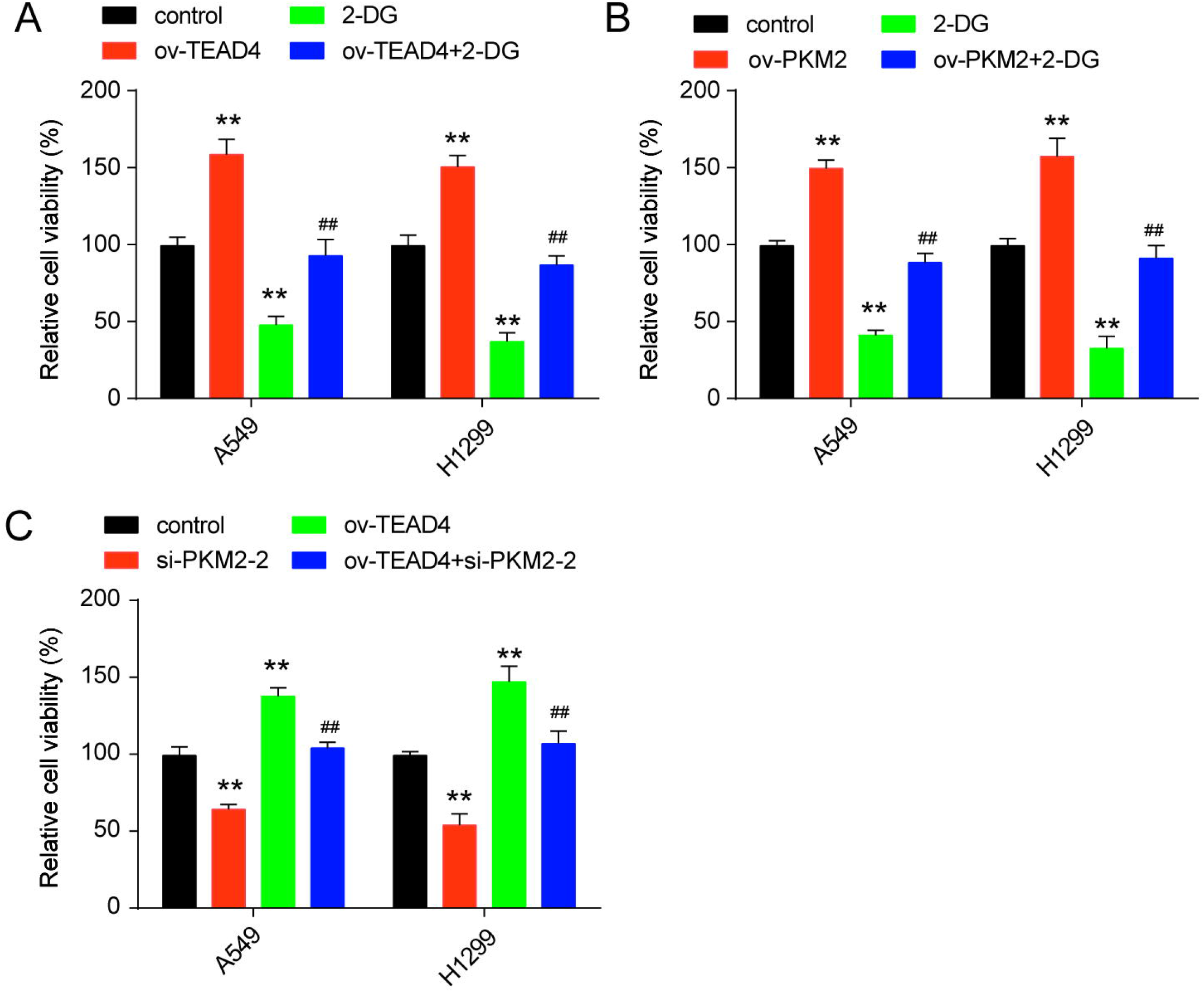
TEAD4/PKM2 axis affects LUAD cells survival through glycolysis. Cell viability were detected by CCK-8 after indicated transfection or 2-DG treatment (10 mM) for 48 h in A549 and NCI☐H1299 cells. (A) 2-DG reversed the acceleration induced by TEAD4 overexpression. (B) 2-DG reversed the acceleration induced by PKM2 overexpression. (C) TEAD4/PKM2 axis promoted LUAD cells survival. \*\**P*<0.01 vs. control, ##*P*<0.01 vs. ov-TEAD4.

## Discussion

More and more evidences show that Hippo pathway is involved in tumor development Yuan et al., 2015. As a vital component of the Hippo pathway in mammals, YAP affects tumorigenesis activity by transcriptional regulation through physical interaction with transcriptional factors TEADs Lin et al., 2017, Zhang et al., 2011. Among four TEAD members, TEAD3 and TEAD4 were reported to be upregulated in LUAD tissues compared with adjacent normal ones Zhang et al., 2018. TEAD4 plays a key role in tumorigenesis and metastasis Zhang et al., 2018. In our study, we also found TEAD4 was upregulated in LUAD tissues and higher TEAD4 level was correlated with poorer overall survival in LUAD. TEAD4 regulated LUAD cells growth via regulating glycolysis through promoting PKM2 transcription. Similarly, TEAD4 has been reported to induce expression of connective tissue growth factor (CTGF) and c☐Myc proteins and then facilitate tumor proliferation of colorectal adenocarcinoma and gastric adenocarcinoma Tang et al., 2018, Zhou et al., 2017. Based on this, we can conclude that TEAD4 was a potential therapeutic target in LUAD.

PKM2 catalyzes the final, irreversible committed step of glycolysis and is highly expressed in cancer cells Zahra et al., 2020. PKM2 plays important role in the genesis and development of tumors, as well as tumoral immune response Sun et al., 2015, Palsson-McDermott et al., 2017. High levels of PKM2 accelerate the capacity of glucose uptake, facilitating cell activation and invasion. During the process of metabolic reprogramming, PKM2 regulates glycolytic pathway in activated immune cells and tumor cells Hou et al., 2020, Sun et al., 2015. Our study indicated that PKM2 was overexpressed in LUAD tissues when compared to noncancerous tissues and was related to the poor prognosis of LUAD. TEAD4 was confirmed to be an upstream regulatory factor of PKM2 in this study. TEAD4/PKM2 axis promoted glucose consumption, lactate production and the extracellular acidification rate. Furthermore, both PKM2 and TEAD4 level in the tumor cells could serve as an independent prognostic factor, which indicates that TEAD4/PKM2 axis is a potential therapeutic pathway to improve anti-cancer efficacy of lung cancer patients.

Except for targeting mutation sites, immune systems and activating signaling, targeting tumor metabolism also have become a hot topic. There are many targets which were regarded as targets for metabolic therapy including key enzymes (GLUT1, HK2 etc.) and product (lactic acid, ROS etc.) Xia et al., 2020. PKM2 was reported to promote the transactivation of HIF-1α via assembling a compound of p300, PHD3 and HIF-1α Luo and Semenza, 2011, increasing expression of genes related to glycolysis in both tumor cells and macrophages Luo et al., 2011, Palsson-McDermott et al., 2015. In this study, we found that TEAD4 and PKM2 was implicated in LUAD cells glycolytic. TEAD4/PKM2 axis altered the reporter activity of HIF-1α and upregulated HIF-1α-targeted glycolytic genes GLUT1 and HK2. Given this association, we can figure out the molecular mechanisms of TEAD4/PKM2 axis on glycolytic.

Summing up, our findings firstly describe that TEAD4 participate in glycolytic of LUAD and PKM2 was a downstream effector of TEAD4. Patients with high expression of TEAD4 or PKM2 in tumor cells have a poorer prognosis. Targeting energy metabolism, especially TEAD4/PKM2 mediated glycolytic, may be a hotspot to enhance anti-cancer efficacy of LUAD patients.

## Materials and methods

### Human tissue sample

Tumor samples were collected from 20 patients, with lung adenocarcinomas and no co-malignancies at The First People’s Hospital of Zigong City. Written consent from the patients and approval from the Ethics Committee of The First People’s Hospital of Zigong City were obtained. None of these patients had received radiotherapy or chemotherapy prior to surgery. Tissue was flash-frozen in liquid nitrogen before long-term storage at −80°C.

### Cell culture and treatment

Human LUAD cell lines, A549, H1299 and human normal lung epithelial cell line HBE were purchased from ATCC. All cells were tested for mycoplasma contamination before used to ensure that they were mycoplasma-free. All cells were cultured in RPMI-1640 medium (Hyclone, Logan, UT, USA) supplemented with 10% fetal bovine serum (FBS, Clark Bioscience, Houston, Texas, USA) at 37°C and 5% CO2.

The overexpression vector and the small interfering RNAs (siRNAs) against human TEAD4 and PKM2 were acquired from Genepharma Technology (Shanghai, China). Lipofectamine 2000 (Invitrogen, Carlsbad, CA, USA) was applied to transfect plasmids and siRNAs into LUAD cells according to the manufacturer’s instruction. The sequences of the siRNA were listed in as follows: si-TEAD4-1, 3’-GTATGCTCGCTATGAGAAT-5’; si-TEAD4-2, 3’-GGACATCCGCCAAATCTAT-5’; si-PKM2-1, 3’-GAAGGAAAUGAUUAAAUCUGG-5’; si-PKM2-2, 3’-GCAAGAUUGAGAAUCACGAGG-5’; si-NC, 3’-GACGACGTACTGTAGTCCA-5’.

### Total RNA extraction and quantitative real-time PCR

Total RNA was isolated by TRIzol Kit (Omega, Doraville, GA, USA) and quantitated by NanoDrop equipment (Thermo Fisher Scientific, Waltham, MA, USA). The PrimeScript kit (Takara Biotechnology, Dalian, China) was used to reverse transcribed. qPCR was performed using SYBR green PCR master mix (Applied Biosystems, Foster, CA, USA). Internal reference genes were set using GAPDH. The normalized method was performed using 2^−ΔΔct^. The primers were list as follows: TEAD4 sense: 3’-TCCACGAAGGTCTGCTCTTT-5’ and anti-sense: 3’-GTGCTTGAGCTTGTGGATGA-5’; PKM2sense: 3’-GCACACCGTATTCAGCTCTG-5’ and anti-sense: 3’-TCCAGGAATGTGTCAGCCAT-5’; GAPDH sense: 3’-TGTGGGCATCAATGGATTTGG-5’ and anti-sense: 3’-ACACCATGTATTCCGGGTCAAT-5’.

### Western blot analysis

Total proteins were extracted using Radio Immunoprecipitation Assay (RIPA) buffer (Beyotime, Shanghai, China) and protein concentration was measured by the BCA Protein Assay Kit (Beyotime, Shanghai, China). Samples were loaded onto 12% sodium dodecyl sulfate-polyacrylamide gel electrophoresis and electrophoretically transferred to polyvinylidene fluoride membranes. Then, the membranes were placed in 5% nonfat milk for blockage and probed with the primary antibodies against TEAD4 (ab151274, Abcam, Cambridge, MA, USA), PKM2 (ab137852, Abcam), GAPDH (ab8245, Abcam), HIF-1α (ab179483, Abcam), HK2 (ab209847, Abcam), GLUT1 (ab195021, Abcam) at 4°C for overnight. The membranes were subsequently subjected to 2 h of incubation with the appropriate corresponding horseradish peroxidase-conjugated secondary antibody (Abcam). Immunoblots were visualized by enhanced chemiluminescence (ECL kit, Beyotime, Shanghai, China). The Western blot bands were quantified by ImageJ.

### Luciferase reporter assay

PKM2 promoter region was constructed into pGL3-based vectors. To determine the effect of TEAD4 on PKM2 promoter, pGL3-based construct containing PKM2 promoter sequences plus renilla luciferase reporter plasmid was individually transfected into A549 and H1299 cells. 48 h after transfection, firefly and renilla luciferase activity were measured by a dual-luciferase reporter assay system (Promega, Madison, WI, USA). The ratio of firefly luciferase to renilla activity was calculated for each sample. To determine the effect of TEAD4/PKM2 axis on reporter activity of HIF-1α, HIF-1 luciferase reporter plasmids (YEASEN, Shanghai, China), which contain multiple HIF-1 binding sites, were co-transfected with TEAD4 overexpression plasmids or si-PKM2-2. 48 h after transfection, firefly and renilla luciferase activity were measured by a dual-luciferase reporter assay system.

### Glycolysis Process Assay

Glycolysis was investigated according to glucose consumption, lactate production and the extracellular acidification rate (ECAR). After transfection for 48 h, the culture medium was collected. The glucose uptake and lactate production were detected using Glucose Assay Kit (Rsbio, Shanghai, China) and Lactate Assay Kit (Rsbio). For the glycolytic function assay, transfected cells were seeded in a XF 24-well plate at a density of 1 × 10^4^ per well overnight and then serum starvation for 24 h. ECAR was detected using Seahorse XF Glycolysis Stress Test kit (Agilent Technologies, Santa Clara, CA, USA) according to the manufacturer’s handbook. The basal ECAR was measured under the basal condition, followed by the sequential addition to each well of glucose (10 mM), oligomycin (2 mM) and 2-deoxyglucose (100 mM).

### Cell survival

Cell survival was measured by the Cell Counting Kit-8 (Beyotime, Shanghai, China). Briefly, 1 × 10^4^ transfected cells were seeded onto 96-well plates per well. After 2-DG treatment for 48 h, 10 μL CCK-8 solution was added and incubated at 37°C for 2 h. The optical density at 450 nm (OD450) was measured for each sample.

### Statistical Analysis

The results were displayed as mean ± standard deviation (SD). Two-tailed Student’s t-test and one-way analysis of variance (ANOVA) followed by Tukey’s test were used to compare the differences in two groups and multiple groups. Kaplan–Meier survival curve was analyzed with log-rank test. P < 0.05 was considered statistically significant.

## Data availability

No publicly available datasets were used for the studies presented in this article.

## Author contributions statement

Y.H.: methodology; investigation, formal analysis, project administration, writing-original draft preparation, visualisation;

H.M.: methodology; investigation, validation, writing-original draft preparation;

Z.D.: conceptualisation, methodology, supervision, writing-review and editing.

